# Decoding transcriptional identity during Neuron-Astroglia Cell Fate driven by RAR-specific agonists

**DOI:** 10.1101/2024.12.23.630055

**Authors:** Ariel Galindo-Albarrán, Aysis Koshy, Maria Grazia Mendoza-Ferri, Marco Antonio Mendoza-Parra

## Abstract

How cells respond to different signals leading to defined lineages is an open question to understand physiological differentiation leading to the formation of organs and tissues. Among the various morphogens, retinoic acid signaling, via the RXR/RAR nuclear receptors activation, is a key morphogen of nervous system development and brain homeostasis. Here we analyze gene expression in ∼80,000 cells covering 16 days of monolayer mouse stem cell differentiation driven by the pan-RAR agonist all-trans retinoic acid, the RARα agonist BMS753 or the activation of both RARβ and RARγ receptors (BMS641+BMS961). Furthermore, we have elucidated the role of these retinoids for driving nervous tissue formation within 90 days of brain organoid cultures, by analyzing > 8,000 distinct spatial regions over 28 brain organoids.

Despite a delayed progression in BMS641+BMS961, RAR-specific agonists led to a variety of neuronal subtypes, astrocytes and oligodendrocyte precursors. Spatially-resolved transcriptomics performed in organoids revealed spatially distinct RAR isotype expression leading to specialization signatures associated to matured tissues, including a variety of neuronal subtypes, retina-like tissue structure signatures and even the presence of microglia.

## Introduction

How cells respond to different signals to develop along defined cell lineages is an open question to understand physiological cell differentiation leading to the formation of organs and tissues, but also to unveil the mechanisms that govern aberrant cell transformation leading to disease. Among the various morphogens, all-trans retinoic acid (ATRA), a vitamin A metabolite, plays a key role in vertebrate embryogenesis, and notably in the development of the nervous system ^1^. ATRA has been shown to act via its interaction with its cognate retinoic acid receptor targets, RARα, RARβ and RARγ, encoded by three different genes and expressed in addition through various isoforms ^2^. During mouse development, all three RAR receptors present spatially distinct expression signatures, indicative for specialized functions (reviewed in ^1^), and such distinct spatial signatures are also retrieved in the adult brain ^2,3^.

Previous studies, including ours, focused on the power of retinoids action for driving cell differentiation in vitro, demonstrated that the use of specific RAR agonists could lead to a variety of cell types ^4–6^. P19 embryonic stem cells treated with the RARα specific agonist BMS753 were shown to give rise to neuronal precursors in a period of 48 hours, but they not progress in differentiation when treated with the RARβ-specific agonist BMS641 or the RARγ agonist BMS961 ^6^. Furthermore, we have previously reported that in 10 days of culture conditions both the RARα agonist BMS753 and the combined use of the RARβ and the RARγ agonists (BMS641+BMS961) lead to a complex differentiation process generating a variety of neuronal subtypes, but also oligodendrocyte precursors and GFAP (+) astrocyte cells ^7^. The analysis of the RAR-driven gene programs revealed that such synergistic action is the consequence of the hijacking of RARα-controlled programs, in agreement with previous studies describing a redundancy for the activation of certain genes by distinct RARs, and notably on Rar-null mutant lines ^5,8,9^.

Concerned by the fact that RARs are known to present anatomically distinct expression signatures both in the embryo but also in the adult brain, we aimed in this study to address the capacity of both the RARα specific agonist BMS753 and the synergistic action of the RARβ and RARγ agonists (BMS461+BMS961) to generate distinct cell types during neuroectodermal differentiation, but also to evaluate their potential spatially distinct expression in *in vitro* 3-dimentional brain organoid cultures. We have first revealed cell types heterogeneity generated during 16 days of mouse stem cell cultures in presence of either of the aforementioned ligand conditions, thanks to the used of single-cell transcriptomics strategy. Then we have setup mouse brain organoid cultures (3 months) in presence of either vitamin A, ATRA, BMS753, or the combination of BMS461+BMS961, to further enhance our understanding of the role of distinct RARs for generating distinct cell-types structured within distinct spatial regions.

## Results

### Synergistic activation of RARβ and RARγ receptors give rise to a variety of neuronal subtypes in mouse embryonic stem cells

With the aim to address potential cell differentiation differences driven by the action of either the pan-RAR agonist All-trans retinoic acid or the use of RAR specific agonists, we have cultured mouse embryonic stem cells in monolayer conditions and treated them with either the pan agonist All-trans retinoic acid (ATRA), the RARα specific agonist BMS753, or the combination of the RARβ and RARγ agonists; BMS641+BMS961. After 8 days of continuous treatment, cells were exposed for other 8 days to neurobasal medium presenting the B27 supplement but devoid of Vitamin A, to promote cell specialization (**Figure 1A**). In aggreement with our previous observations ^7^, the use of either of these conditions allowed to obtain mature neuronal cell differentiation, as confirmed by immunofluorescence using the mature neuronal marker MAP2 (**Figure 1B**).

**Figure 1.**
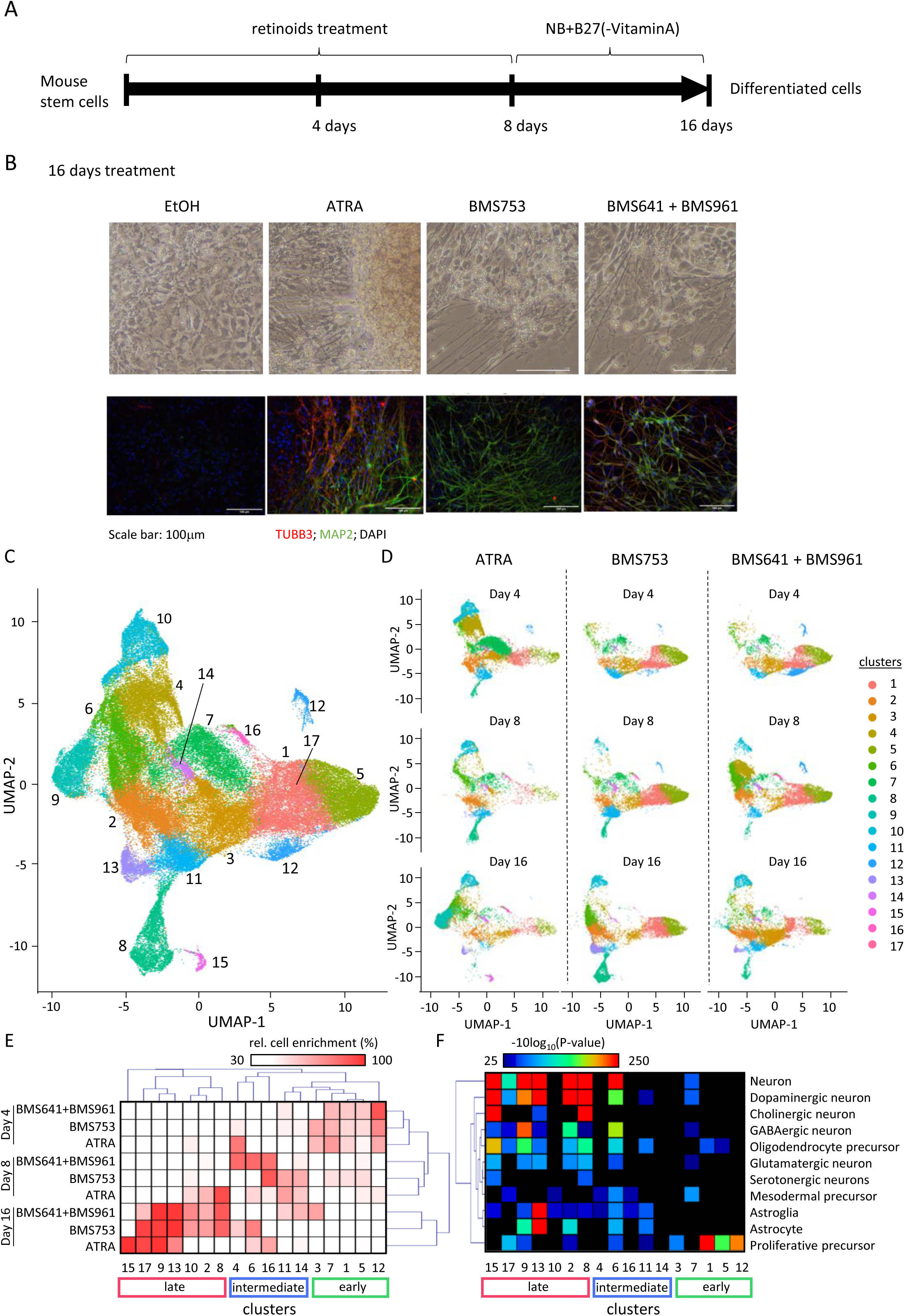
Transcriptome at single cell level reveals retinoids-driven stem cells specialization differences in a temporal dependency. **(A)** Scheme illustrating the mouse stem cell differentiation strategy, where cells were treated with different retinoid agonists (*all-trans* retinoic acid: ATRA, RAR-alpha agonist BMS753, or combination of the RAR-beta agonist BMS641 and the RAR-gamma agonist BMS961) during 8 days, then exposed to maturation media (Neurobasal medium: NB, supplemented with B27 devoid of Vitamin A). Cells were collected at 4, 8 or 16 days for single cell transcriptome assays. **(B)** Representative micrographs assessed in either bright-field (top panels) or immunofluorescence (bottom panels) from cells treated with Ethanol (EtOH), ATRA or the combination of BMS641 and BMS961 analyzed after 16 days of differentiation. Cells were stained for the neuronal precursor marker TUBB3 (red), and the marker for mature neurons MAP2 (green). Nuclei were stained with DAPI (blue). **(C)** Uniform Manifold Approximation and Projection (UMAP) for all three conditions (ATRA, BMS753, BMS641+BMS961) assessed at three time-points (4,8, 16 days). Color-code correspond to 17 non supervised clusters **(D)** UMAP illustrated in (C) stratified per treatment and time-point. **(E)** Heatmap representation of the relative number of cells per cluster and normalized across time-points per treatment. UMAP clusters and conditions are clustered by “Euclidean” distance. **(F)** Cell-type annotations per corresponding UMAP cluster assessed by hypergeometric probability assay. Colored boxes in (E) and (F) make reference to the group of clusters retrieved associated to either early, intermediate or late stages during differentiation.

While in our previous study we have demonstrated that the synergistic activation of the RARβ+RARγ receptors recapitulate neuronal cell differentiation like in the case of the use of the RARα agonist, in this study we focus our attention to the potential differences in neuronal subtypes that can be generated by using different RAR receptors. To address this question we have collected samples during retinoids treatment (4 days), before adding the neurobasal medium (8 days), and at the end of the 16 days of cell differentiation. Single-cell transcriptomics assays performed in 79.596 cells across all samples and treatment conditions were analyzed together by using a Uniform Manifold Approximation and Projection for dimension reduction (UMAP) revealing up to 17 distinct cell clusters (**Figure 1C**). Stratification of this global UMAP visualization by treatments and time-points revealed differences between treatment conditions. Notably, cluster 8 has been retrieved in early stages of ATRA treatment (d4: 4%, d8: 19% of cells) and in late stages of BMS641+BMS961(d8: 2%, d16: 7% of cells), but it is preferentially enriched in BMS753 treatment (d8: 2%, d16: 61% of cells) (**Figure 1D; Supplementary Table S1**). Furthermore, cluster 12 is mainly seen in BMS641+BMS961 treatment (d4: 69% of cells); cluster 4 appears preferentially in ATRA at day 4 (50% of cells) and in BMS641+BMS961 at day 8 (34% of cells); and cluster 3 is retrieved in BMS641+BMS961 at day 16 (32% cells) and in ATRA treatment at day 4 (20% of cells) (**Figure 1D & Supplementary Table S1**).

A classification strategy applied to the relative enrichment levels of cells per cluster and adjusted across timepoints allowed to highlight a correlation with the timeframe. Indeed, clusters 3, 7, 1, 5, and 12 were associated to early events (day 4); clusters 4, 6, 16, 11 and 14 to intermediate transitions (day 8); and cluster 15, 17, 9, 13, 10, 2 and 8 to late events (day 16); and this in despite of the treatment in use (**Figure 1E**). Importantly, this analysis also revealed discrepancies between ATRA and the other ligand treatments, including a specific ATRA enrichment in cluster 15, and temporally desynchronized behavior relative to ATRA in clusters 10, 2, 8; 16; 11; indicative of a delayed maturation process driven by the specific ligands relative to the pan-RAR agonist. Furthermore, clear differences between the BMS641+BMS961 and the BMS753 treatment were observed. Cluster 4&6 presented a preferential enrichment in day 8 for BMS641+BMS961 treatment (cl4-d8: ∼81% cells per condition adjusted to timepoint in BMS641+BMS961; cl6-d8: ∼82% in BMS641+BMS961); and cluster 3&11 in day 16 (cl3-d16: ∼62%; cl11-d16: ∼46% of cells in BMS641+BMS961). In contrary, cluster 11 displayed a preferential enrichment in day 8 for BMS753 treatment (d8: 60% BMS753); as well as cluster 17 in day 16 (d16: ∼95% BMS753) (**Figure 1E & Supplementary Table S1**).

A cell-type annotation strategy, based on public gene expression signatures, allowed to clearly state that clusters associated to the earlier timepoint corresponded to proliferative precursors (clusters 3, 7, 1, 5 and 12) (**Figure 1F & Supplementary Table S2**). The intermediate cluster 6 (day 8) presented signatures associated to neurons, including Dopaminergic and GABAergic subtypes, as well as oligodendrocyte precursors and Astrocytes. Finally, clusters associated to day 16 presented enhanced signatures for the aformentioned neuronal subtypes, as well as new others including chrolinergic (cluster 8, 13,15) and serotonergic neurons (cluster 8, 15). Importantly, among the previoulsy preferentially associated clusters to the BMS641+BMS961 treatment, cluster 6 appeared to be enriched in Dopaminergic, and GABAergic neuronal subtypes, as well as signatures associated to Oligodendrocyte precursors and astroglia (**Figure 1F**). In contrary, cluster 11 – preferentially enriched to the BMS753 treatment – presented weak signatures associated to Oligodendrocyte precursors and astroglia, while cluster 17 presented signatures associated to Dopaminergic neuron subtype (**Figure 1F**).

Overall, mouse embryonic stem cells treated with the combination of the RARβ and RARγ agonists recapitulated the differentiation potential driven by either the pan-agonist ATRA or the RARα agonist BMS753; with some differences on the grounds of the obtained neuronal subtypes. Despite these differences, several of the identified clusters during this analysis appeared associated to multiple cell-types, suggesting that a further celltype stratification strategy might be required for better resolve such potential differences.

### disentangled RARβ+RARγ driven trajectories during differentiation reveals a delayed progression relative to RARα activation

To further stratify the cell clusters obtained by UMAP classification, we have taken advantage of STREAM, a pseudo-time stratification methodology ^10^. At first, we have stratified at once all cells collected in all three timepoints and issued from all treatment conditions. This analysis gave rise to 14 pseudo-time trajectories, from which, pseudotime S2 corresponded to the main root composed of cells harboring a strong transcriptomics signature similarity (**Figure 2A**). This main root gave rise to 4 intermediate branches (S4, S12, S6, S5), leading in a second time to other 9 terminal branches (S1, S8, S7, S3, S14, S13, S11, S9, S10) (**Figure 2A**). Importantly, cells issued from the BMS753 treatment were preferentially enriched in the terminal branches S13 (d16: 79%), S7 (d16: 86%) and in the intermediate branche S4 (d16: 53%) (**Figure 2B,C & Supplementary Table S3A**). Terminal branches S9 (d4: 96%), S3 (d4: 59%; d8: 24%), S14 (d8: 57%), S8 (d16:85%) and the intermediate branche S5 (d4: 81%) were enriched for cells issued from ATRA treatment. Finally, the BMS641+BMS961 treatment appeared to contribute preferentially to only two intermediate branches (S1_d8: 45% and S6_d16: 68%), or retreaved together with BMS753 in multiple other branches (S2 (d4:27%;d8: 18%), S4 (d16:33%), S12 (d16:18%), S11 (d16:17%), S10 (d16:20%) **(Figure 2B,C & Supplementary Table S3A**)) suggesting that this treatment is redundant with the outcome of the BMS753, as demonstrated in our previous study ^7^.

**Figure 2.**
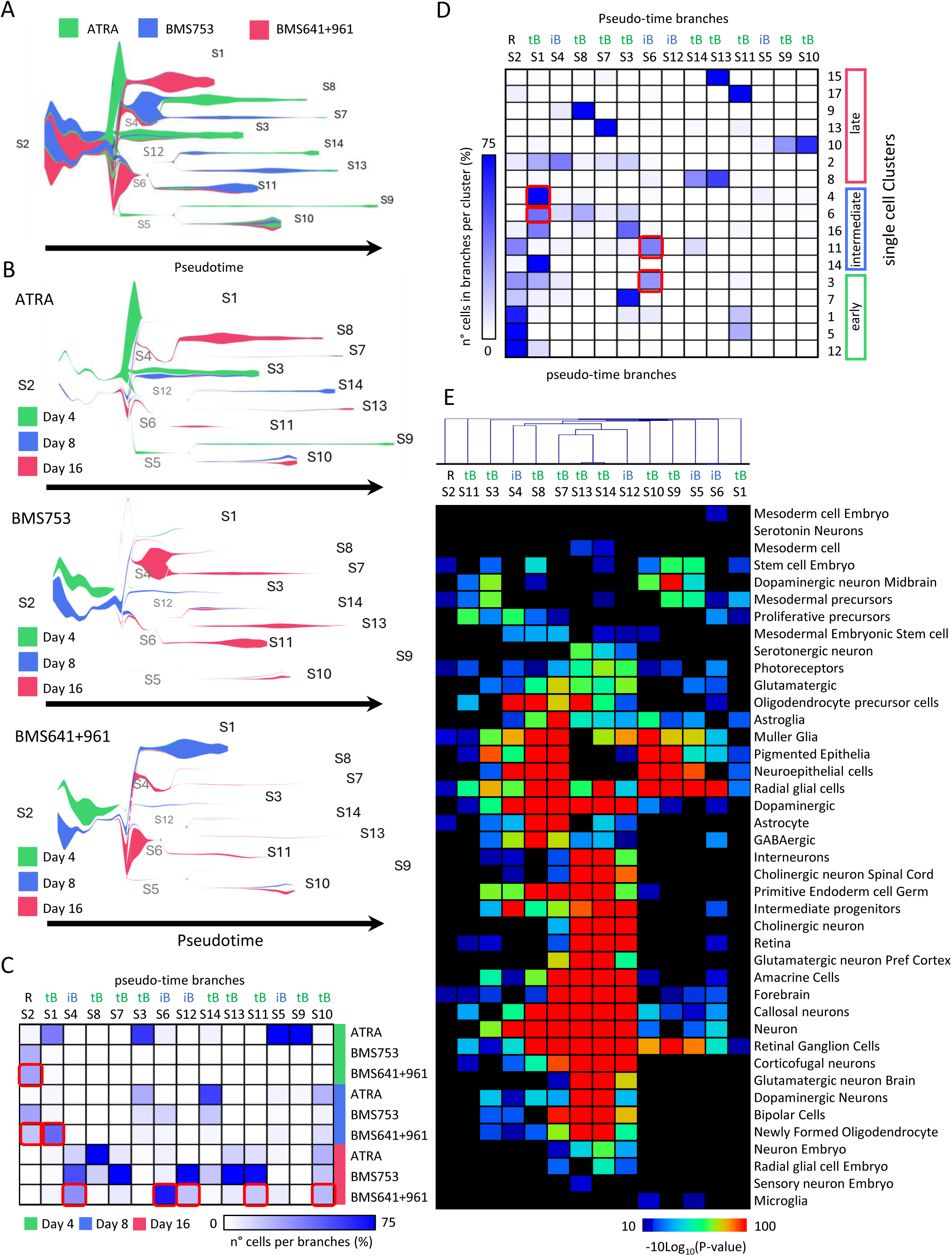
Pseudo-temporal cells stratification reveals a delayed cell specialization progression in RARβ+RARγ-driven trajectories relative to the RARα activation. **(A)** Pseudo-time stratification of single-cell transcriptomics data assessed in all three treatments. **(B)** All 14 branches assessed in (A) are displayed per treatment and colored on the grounds of their associated time-points. **(C)** Fraction of cells per pseudo-time branches (in percent) displayed in the context of their corresponding treatment and the collected time-point. **(D)** Fraction of cells in pseudo-time branches per UMAP clusters displayed in Figure 1. In (D) and (E) red square boxes correspond to fraction of cells associated to the BMS641+BMS961 treatment. **(E)** Heatmap displaying cell-type annotation (hypergeometric probability gene set enrichment assay) performed per pseudo-time branches (clustered by average dot product distances).

A direct comparison between clusters obtained by the UMAP classification and the pseudo-time progression provided by STREAM allowed to further stratify their composition. For instance, cluster 6, initially described as BMS641+BMS961 specific, presented a strong overlap with pseudotime branche S1 (42%), S4 (11%), S8 (25%) and S3 (13%) (**Figure 2D & Supplementary Table S3B**). Among all these four pseudo branches, only S1 and S4 has been associated to BMS641+BMS961, while S8 and S3 to BMS753 and/or ATRA treatment (**Figure 2C**). Similarly, cluster 11 (associated to BMS753) has been enriched in pseudotime root S2 (36%), the intermediate branche S6 (37%) and the terminal branche S14 (11%). Or, as illustrated in **figure 2C**, the branche S6 has been enriched in cells treated with BMS641+BMS961 and branche S14 with ATRA.

In the other hand, events taking place in the late phase of differentiation appeared more specific between cluster and pseudo-time branches. For instance, the terminal branche S11 and cluster 17 appeared specifically enriched for cells issued trom the BMS753 treatment; branche S13 corresponded to clusters 8 &15; S14 is mainly enriched in cluster 8, S8 in cluster 9, S7 in cluster 13, and pseudo-time branches S9 &S10 in cluster 10 (**Figure 2D**).

Cell-type annotation assay revealed that the pseudo-time terminal branches S13 & S14 (mainly associated to ATRA and BMS753 treatment) are strongly enriched for gene expression signatures related to forebrain development, amacrine cells, Retina, retinal ganglion cells, callosal neurons, corticofugal neurons, as well as to cholinergic, glutamatergic or dopaminergic neuronal markers (**Figure 2E**). Terminal branches S8, S7 (mainly associated to ATRA and BMS753 treatment) presented signatures related to astroglial cells including radial glial cell, astrocyte, oligodendrocyte precursors in addition to Muller glial, pigmented epithelia or neuroepithelial cell markers (**Figure 2E**). Several of these markers were also retrieved in branches S9 (ATRA-specific) but also in S10, indicative for the presence of such type of especialized cells under BMS641+BMS961 treatment. Furthermore, the other pseudo-time branches associated to BMS641+BMS961 treatment were enriched in gene expression signatures associated to radila glial cells (S6), Intermediate progenitors, neuroeptihelial cells, oligodendrocyte/proliferative precurors (S4), arguing for a less differentiated status than those observed in branches S13,S14, associated to ATRA or BMS753 treatment (**Figure 2E**).

### Repressive/active epigenetic signature transitions at the promoter level controls cell differentiation progression in RARα or RARβ+RARγ activation

Considering the differences observed in single-cell transcriptomics assays performed between BMS753 and BMS641+BMS961, we aimed at evaluating for potential epigenetic signatures retrieved at promoter regions (5kb around Transcript start site) able to confirm such observations. To do so, we have performed Cleavage Under Targets and Tagmentation (Cut&tag ^11^) assays after 16 days of BMS753 or BMS641+ BMS961 driven cell differentiation, targeting the repressive histone modification mark H3K27me3, the active promoter markers H3K27ac and H3K4me3, as well as the RNA polymerase II (RNAP2). The obtained chromatin enrichment signatures were integrated for revealing promoter co-occurring combinatorial events. As illustrated in **Figure 3A**, a total of 10 chromatin co-occuring events were retrieved, which were associated to functional promoter status as following: Promoters being in a transcribing active status (presenting both the H3K27ac and the H3K4me3 markers and if possible also traces of RNAP2, or RNAP2&H3K27ac); promoters presenting an active enhancer status (presenting only H3K27ac); promoters being repressed (enriched in H3K27me3), or simply being devoid of any of these markers and thus considered as inactive (**Figure 3A**).

**Figure 3.**
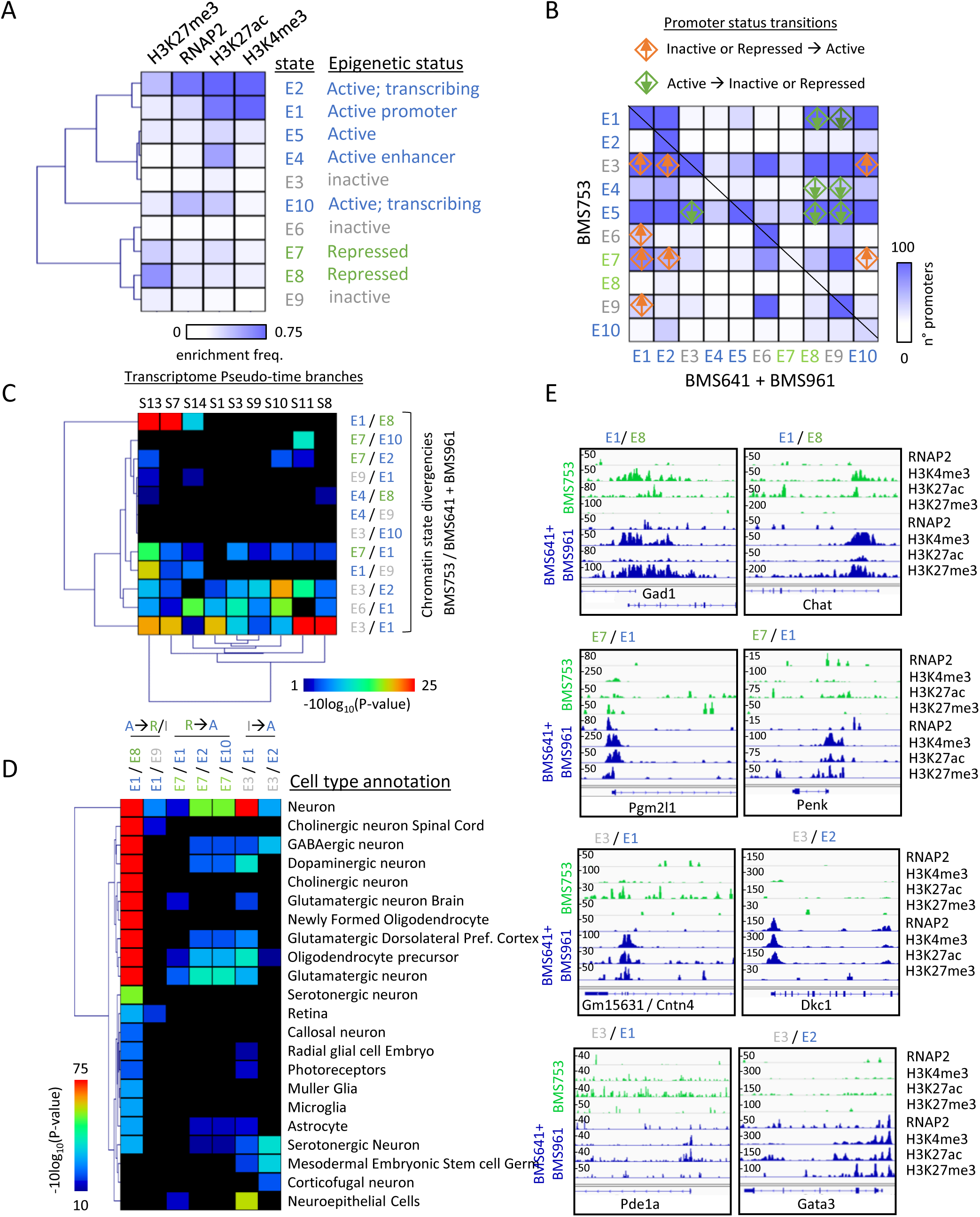
Epigenetic signature transitions at the promoter level controls cell differentiation progression in RARα or RARβ+RARγ activation. **(A)** Chromatin state analysis performed on chromatin enrichment signatures (Cut&tag) assessed for the repressive mark H3K27me3, RNA polymerase II (RNAP2) and the active promoter markers H3K27ac and H3K4me3; all collected in mouse stem cells differentiated during 16 days in presence of either the RARα ligand BMS753, or the combination of the RARβ and RARγ agonists (BMS641+BMS961). Ten different combinatorial states were retrieved (ChromHMM), which were classified based on their promoter functional roles. **(B)** Comparison between the promoter chromatin status assessed in samples issued from the BMS753 and those issued from the BMS641+BMS961. The illustrated heatmap displays the number of promoters sharing the indicated chromatin state for either conditions. Diamond symbols indicate promoters presenting an inactive or repressed state in BMS753 and being active in BMS641+BMS961 (orange diamond), or the opposite (green diamond). **(C)** Heatmap comparing the promoter chromatin state transitions between BMS753 and BMS641+BMS961 and the corresponding gene expression associated to the pseudo-time branches displayed in Figure 2. **(D)** Cell type annotation (hypergeometric probability gene set enrichment assay) of the genes associated to the most relevant promoter chromatin state transitions observed in (C). **(E)** Example of promoter chromatin state transitions between BMS753 and BMS641+BMS961 conditions. Promoter enrichment levels were assessed in a window of 5kb around the transcription start site.

Once promoter satus are characterized, we have classified signatures found in either of the retinoids treatment to retrieve for potential changes. Indeed, while the majority of genes are presenting a similar histone modification signatures within their promoters, we have identified several genes passing from an inactive/repressed status in BMS753 into a trancscriptionally active status in presence of BMS641+BMS961 (E3/E1:312 genes; E3/E2:192 genes; E3/E10:108 genes; E6/E1: 35 genes; E7/E1: 82 genes; E7/E2: 49 genes; E7/E10: 18 genes; E9/E1: 29 genes), as well as others doing the oposite (E1/E8: 517genes; E1/E9: 72 genes; E4/E8: 23 genes; E4/E9: 41 genes; E5/E3: 127 genes; E5/E8: 1117 genes; E5/E9: 1216 genes) (**Figure 3B**).

The comparison of this information with the pseudo-time branches described in **Figure 2** led to the identification of a strong enrichment of promoters passing from an active status in BMS753 into a repressive/inactive status in the presence of BMS641+BMS961 in branches S13, S7 (E1/E8 transitions). Furthermore, branches S11, S8, S1 were preferentially enriched for a promoter activation transition in the combination of the RARb+RARg agonists (E3/E1); confirming the results obtained from the sigle-cell transcriptomics (**Figure 3C**).

Finally, a cell type annotation analysis confirmed that active promoters retrieved in the BMS753 condition but repressed in the BMS641+BMS961 treatment (E1/E8) were preferentially associated to genes related to a variety of neuronal subtypes; while active promoters in BMS641+BMS961 treatment but repressed in BMS753 failed to present a distinct signature, with the exception that were enriched for markers associated to neuroepithelial cells (E3/E1), indicative for a less differentiated state than that observed in BMS753 (**Figure 3D**).

Examples of such promoter status transitions are illustrated in **Figure 3E**, where genes like Gad1 (Glutamate Decarboxylase 1; known as marker for GABAergic neurons), or Chat (Choline O-Acetyltransferase; known to catalyze the biosynthesis of the neurotransmitter acethylcholine) were found to present an active promoter status in BMS753, but rather repressed in BMS641+BMS961 treatment (E1/E8 transition). In contrary, A repressive to active transcriptional behavior (E7/E1) has been observed in the promoter associated to genes like Pgm2l1 (Phosphoglucomutase 2 Like 1; coding for an enzyme strongly expressed in the brain, accounting for the elevated levels of glucose-1,6-biphosphate), or the gene coding for the Proenkephalin protein (Penk), a neuropeptide that plays a role in the perception of pain and response to stress, and which its expression under the control of RARγ in DARPP32-positive striatal-like medium spiny neurons ^12^ (**Figure 3E**). Similarly, genes like Cntn4 (Contactin4, an axon associated cell adhesion component, playing a role in neuronal netwok formation and plasticity), Dkc1 (Dyskerin Pseudouridine Synthase 1, a protein involved in the telomerase activity), Pde1a (Phosphodiesterase 1A, a protein known to be expressed in neurons where it hydrolizes cGMP into cAMP) or Gata3 (transcription factor known to play a major role in the development of serotonergic neurons ^13^), were shown to present an inactive promoter status in presence of the BMS753 treatment but present a rather active state in presence of the BMS641+BMS961 (**Figure 3E**).

Overall, these disentangle analyses performed either in the single cell transcriptomics data with the help of pseudo-time inference strategies, or the assessment of promoter status by the capturing of relevant histone modification signatures, confirm the capacity of the BMS753 and the combination of the BMS641+BMS961 RAR ligands to drive the emergence of a variety of cell types during cell differentiation, and notably a delayed progression in differentiation in the case of the synergistic activation of the RARβ and RARγ receptors.

### Long-term brain organoid cultures in presence of RAR-specific agonists provide distinct nervous tissue differentiation signatures

The aforementioned differentiation assays performed in 2-dimensional monolayer cultures demonstrated (i) a delayed progression towards maturation in presence of the combination of the RARβ and RARγ agonists BMS641+BMS961 relative to the use of the RARα agonist BMS753; and (ii) subtle differences in neuronal subtypes generated in these two conditions. Considering the major relevance of the retinoids action during brain development ^1,14^, and notably the use of vitamin A (the natural precursor of the pan-RAR agonist, the all-trans retinoic acid) for maturation of cerebral organoids in *in vitro* 3-dimensional cultures ^15^, we have envisioned that the use of RAR-specific agonists could be considered as an option for generating specialized cerebral organoids. To address this hypothesis, we have followed the previous protocol described by M. Lancaster for generating mouse brain organoids ^14^, with the slight difference that in day 9 we have transferred the matrigel embeeded organoids into flasks instead of bioreactors (**Figure 4A**). Furthermore, to address the potential performance of retinoids action for organoids maturation we have complemeted the maturation media (Day 9) with either Vitamin A, All-trans retinoic acid (ATRA), the RARα agonist BMS753, or the combination of the RARβ and RARγ agonists (BMS641+BMS961) (**Figure 4A & Supplementary Figure S1A**).

**Figure 4.**
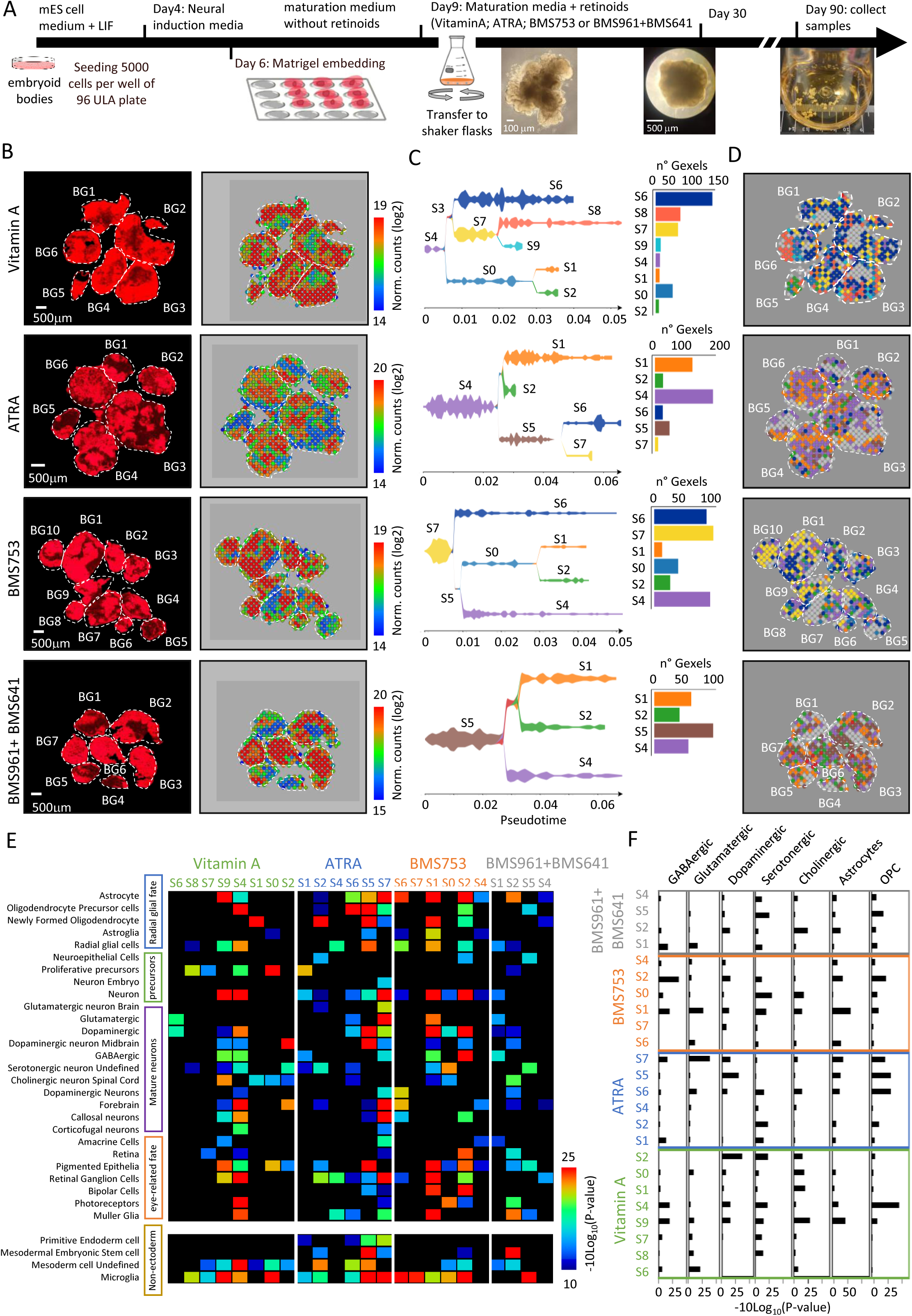
Long-term brain organoid cultures in presence of RAR-specific agonists provide distinct nervous tissue differentiation signatures. **(A)** Scheme illustrating the strategy in use for generating mouse brain organoids. After 90 days of culture, multiple brain organoids are collected for spatial transcriptomics assay **(B)** Left panels: Microscopy image from a double-barcoded DNA array slide presenting a cryosection of multiple brain organoids per treatment, in which captured cDNA is visualized by dCTP-Cy3 incorporation. Right panels: Corresponding spatial transcriptomics landscapes assessed with the help of the in-house double-barcoded DNA arrays (Lozachmeur et al. ^18^). Heatmap landscapes correspond to the normalized read counts per Gexel (gene expression element; in analogy to Pixel). BG: Brain organoid. **(C)** Left panels: Gexels stratification per treatment condition with the help of a pseudo-time strategy (STREAM^10^). Right panels: number of Gexels per pseudo-time branches. Spatial localization of the identified branches in (C). Gray Gexels present low normalized read-counts in (B), thus they are not included in the pseudo-time analysis. **(E)** Heatmap displaying gene set enrichment levels for cell/tissue type annotations performed over all pseudo-time branches across all treatments. **(F)** Barplot illustrating the gene set enrichment confidence assessed for neuronal subtype signatures across the various pseudo-time branches issued from the various brain organoid treatments.

Brain organoids were traced over time by RT-qPCR targeting the depletion of pluripotency markers, the transient emergence of neuronal precursor signatures, as well as the establishment of neuronal subtype markers. Among the evaluated pluripotency markers, Oct4 appeared strongly downregulated in both VitaminA and ATRA treatment after 90 days of differentiation. In contrary, the use of either BMS753 or BMS641+BMS961 gave rise to mild detected downregulation, indicative for the presence of pluripotent cells even after 90 days of culture in these two conditions (**Supplementary Figure S1B)**. In the other hand, strong levels of the Klf4 transcript were detected in all conditions and across all time-points, indicative for the presence of neuronal progenitors ^16,17^. Neuronal precursors were confirmed also by the detection of transcripts associated to Ascl1, Nestin and Tuj1; which presented high levels during the first 30 days, but were gradually reduced under all treatment conditions. Furthermore, transcripts associated to the gene Gad67 (GABAergic neurons), Glut1 (Glutamatergic neurons), Th (Tyrosine Hydroxulase, marker for Dopaminergic neurons) and Tph2 (tryptophan hydroxylase, marker for Serotonergic neurons) were detected preferentially under VitaminA treatment **(Supplementary Figure S1B)**. Indeed, ATRA and BMS753 treatment conditions presented barely 2-fold levels of the Tph2 transcript relative to the day 0 control, and completely absent in BMS641+BMS961. Finally, transcripts targeting the astrocyte marker Gfap has been detected in all four treatment conditons treatments and across all timepoints, while the transcript Olig2, specific fo Oligodendrocytes remained undetectable in all treatments **(Supplementary Figure S1B).**

With the aim to perform a more in-depth analysis, after 90 days of culture in presence of retinoids, brain organoids were cryosectioned and tissue sections were deposited in in-house manufactured DNA arrays presenting 2,048 unique molecular positional barcodes and poly-T ends for capturing messanger RNA ^18^. This in-house spatial transcriptomics strategy allowed to capture gene expression signatures across multiple brain organoids cryosectioned together (**Figure 4B**). Gene expression readouts assessed per physical position, herein called as “gexels” (gene expression elements, in analogy to pixels) were stratified in multiple pseudo-time branches with the help of STREAM ^10^. As illustrated in **Figure 4C**, pseudo-time stratification of gexels based on their gene expression levels issued from VitaminA treatment assessed in 6 different brain organoids allowed to identify up to 8 branches. Among them, the branch S6 appeared as the most frequent (∼150 gexels), followed by branches S8 and S7 (∼70 gexels each); all three retrieved in 5 over 6 analysed organoids (**Figure 4D**).

Similar pseudo-time stratification performed over all three other treatment conditions revealed up to 6 branches for ATRA treatment, 7 for BMS753, but only 4 in the case of BMS641+BMS961, and this despite a similar number of analysed brain organoids (**Figure 4B&C**). Indeed, gexels associated to all four branches were retrieved in all 7 analysed brain organoids issued from the BMS641+BMS961 treatment, indicative for an homogeneous differentiation behavior; an aspect that is also retrieved in all other treatments (**Figure 4D**).

Cell-type annotation performed on all identified pseudo-time branches revealed signatures ranging from proliferative precursor states, passing through intermediate progenitors leading to radial glial fate giving rise to astrocytes or driven towards neuronal cell specialization (**Figure 4E & Supplementary Figure S2**). Indeed, traces of proliferative precursor signatures were found enriched in the root S4, and branches S0, S7 & S8 from Vitamin A; S1 from ATRA; and S2 from BMS641+BMS961 treatment (**Figure 4E**). Neuroepithelial cell signatures were found in branches S2 from ATRA treatment; S2 from BMS753; and S5 from BMS641+BMS961 treatment (**Figure 4E**). Radial glial cell signatures were retrieved in S8, S4 & S9 from Vitamin A; S4 & S5 from ATRA treatment, S1, S2 & S6 from BMS753; and S1 & S2 from BMS641+BMS961 treated organoids (**Figure 4E**). Similarly, Astrocyte and oligodendrocyte precursors were enriched in S4 & S9 from Vitamin A; S5, S6 & S7 from ATRA; S1& S2 in BMS753; and S2 & S5 from BMS641+BMS961 treatment; indicative for their cell differentiation with the aforementioned branches (**Figure 4E&F**).

Neuronal signatures were retrieved in the therminal branches S1, S2 & S6 issued from the BMS753 treatment; S4, S9 in Vitamin A; S7 & S5 in ATRA; as well as in S1 from the BMS641+BMS961 treatment (**Figure 4E**). Furthermore, specialized neuronal subtype signatures like those corresponding to GABAergic neurons were found in branches S4& S9 in Vitamin A; S1& S7 in ATRA; S1 & S2 for BMS753; and S1 from BMS641+BMS961 treatment (**Figure 4F**). Similar analysis confirmed enrichment signatures for Glutamatergic, Dopaminergic, Serotonergic and Cholinergic neurons in various pseudo-time branches across all treatments, demonstrating that brain organoid cultures in presence of RAR-specific agonists can lead to well differentiated tissues, presenting a variety of cell-types.

Finally, further functional nervous tissue annotation signatures were enriched in various of the analysed branches; among them the presence of callosal projection neurons (S5 & S7 ATRA; S2 & S6 BMS753; S4 & S9 Vitamin A); bipolar cells (S5 & S7 ATRA; S1 & S2 BMS753); pigmented epithelia (S7 ATRA; S1 & S2 BMS753; S0, S4 & S9 Vitamin A; S5 BMS641+BMS961); retinal ganglion cells (S4, S2, S7 ATRA; S4 BMS641+BMS961; S1, S2, S6 BMS753; S4, S9 Vitamin A); photoreceptors (S0, S2 BMS753; S2 BMS641+BMS961; S4 Vitamin A; S7 ATRA); retina (S2 BMS641+BMS961; S7 ATRA; S2 BMS753; S4, S7 Vitamin A); amacrine cells (S4 BMS753; S1 BMS641+BMS961; S7 ATRA; S9 Vitamin A); or Corticofugal neurons (S7 ATRA; S4 Vitamin A) (**Figure 4E**).

It is also worth to mention that strong microglia signatures were also detected in Vitamin A-treated organoids (S0, S4, S9, S8); in ATRA treatment (S7, S5, S6, S2); in BMS753 (S7, S6, S2, S1); and in a less pronounced manner in BMS641+BMS961 treatment (S5); in aggreement with brain organoid cell differentiation cultures where no means to inhibit the endodermal/mesodermal differentiation pathways are used ^19^.

Overall, this detail analysis of the various tissular regions disected by spatial transcriptomics performed on brain organoid cultures demonstrated that the RAR-specific agonist BMS753 (RARα specific), as well as the synergistic activation of the RARβ and RARγ receptors (BMS641+BMS961) could perform as well as the pan RAR-agonist ATRA, or the lagely used Vitamin A supplement.

### Reconstruction of the RAR-driven gene regulatory programs involved in cell specialization during brain organoids progression

Stratification of the various gexels composing the spatial transcriptomic maps assessed under the various retinoids treatment revealed a variety of gene expression signatures associated to distinct differentiated cell types. Considering that these processes are expected to be driven by the action of the added retinoids, we aimed at evaluating the expression of each of the RAR receptors. As illustrated in **Figure 5**, brain organoids collected at 90 days of BMS753 treatment presented expression signatures for all three *Rara*, *Rarb* and *Rarg* receptors (**Figure 5A**). Such expression signatures were observed in nearly all nine analysed brain organoids, covering up ∼60 gexels for *Rarg*, ∼75 gexels for *Rara*, and ∼60 gexels for *Rarb* (**Figure 5B**). Similar observation was made for these receptors in brain organoids treated with BMS961+BMS641, where ∼80 gexels were associated with upregulated transcripts for *Rara*, ∼55 gexels for *Rarb*, and ∼70 gexels for *Rarg* (**Figure 5A&B**). RAR expression across the analyzed brain organoids represented ∼ 60% of the total Gexels; with less than 5% of them corresponding to presence of all three RARs in the same spatial regions (**Figure 5C**). Barely ∼25% of Gexels over-expressed two over three RAR receptors, and ∼30% corresponded to Gexels presenting only one of the RARs being over-expressed (**Figure 5C**).

**Figure 5.**
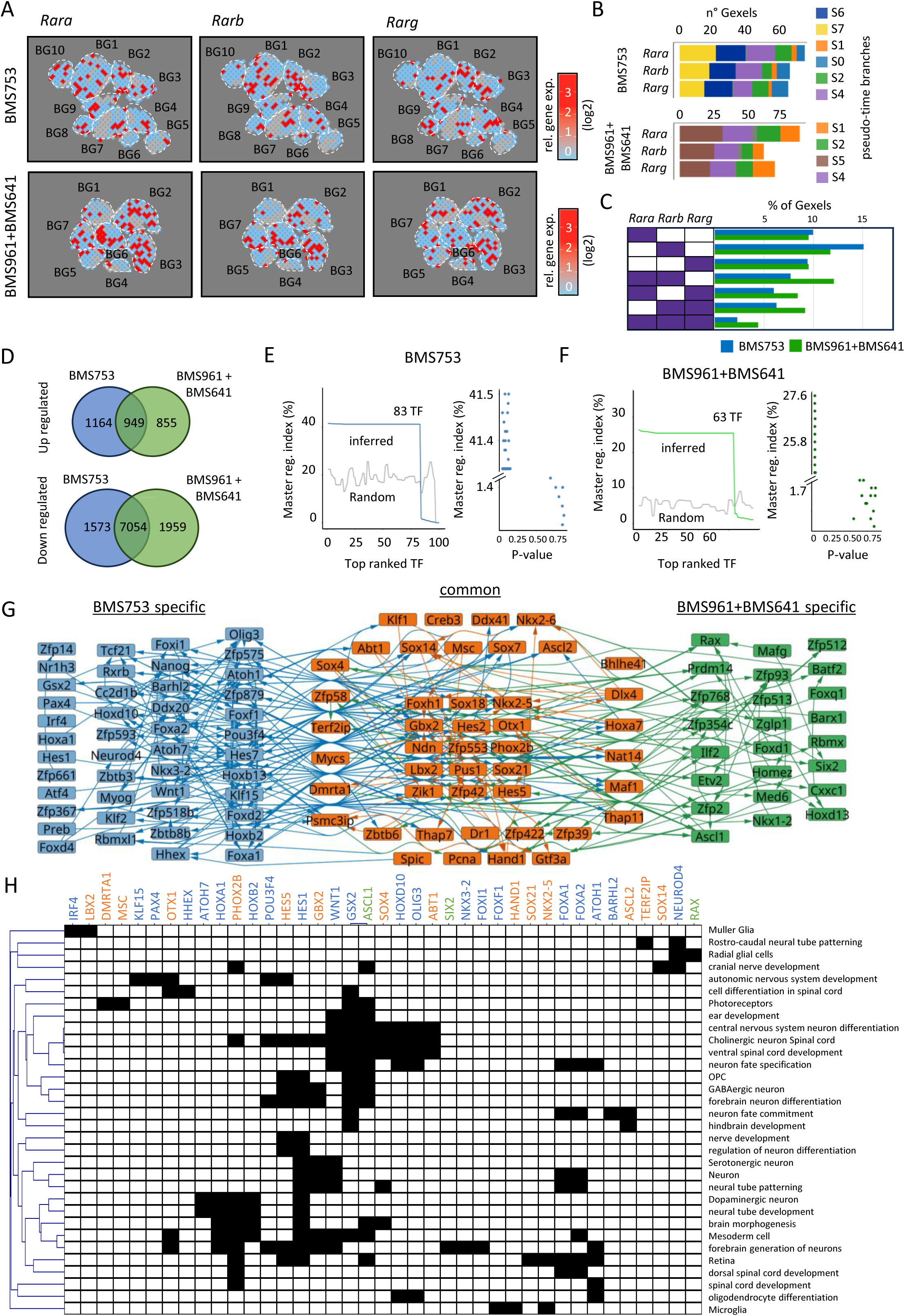
Reconstruction of the RAR-driven gene regulatory programs involved in cell specialization during brain organoids progression. **(A)** Gexels presenting high expression levels of the genes Rara, Rarb or Rarg in brain organoids under the retinoid agonist BMS753 (top panels) or BMS961+641 (low panels). **(B)** Gexel quantification for each Rar gene and the correspondences to pseudo-time branches displayed in Figure 4. **(C)** Fraction of gexels expressing all three RARs, two over three or only one of them, in both treatment conditions. **(D)** Venn diagrams comparing highly expressed genes (log2 fold-change > 1 respect to the logarithmic media) between the treatments MBS753 and BMS961+BMS641. **(D)** Left panel: Ranking of transcription factors in BMS753 treatment, based on their capacity to control downstream factors in a gene regulatory network (GRN) reconstructed from the differentially expressed genes displayed in (D). The master regulatory index (MRI) corresponds to the fraction of genes that can be controlled by a given TF within the GRN (TETRAMER^20^). MRI inferred to the top ranked TFs are displayed in blue solid line. In gray, a randomized GRN of the same size has been used for evaluating the MRI confidence. Right panel: Confidence associated to the TF computed from the randomized GRN. **(F)** Same as (E) for the differentially expressed genes assessed from brain organoids under the BMS961+BMS641 treatment. **(G)** Gene co-regulatory network of the top ranked master TFs obtained in (E) and (F). TFs specific to the MBS753 treatment are displayed in blue, those specific to BMS961+641 in green and those in common for both treatments in orange. The network was organized from outside to inside by the topological parameter “betweenness centrality” (most incoming connections at the network center). **(H)** Cell/tissue type annotations associated to the most relevant TFs retrieved in the gene co-regulatory network.

Gexels presenting over-expressed transcripts for at least one of the RAR receptors in either of treatments were compared on the grounds of their differentially expressed genes. This comparison revealed 949 common upregulated genes, 1,164 upregulated genes specific to the BMS753 treatment and 855 specific to BMS961+BMS641 (**Figure 5D**). Surprisingly, > 7,000 genes were found commonly downregulated, where only 1,573 genes were specific to BMS753 and 1959 to BMS961+BMS641; indicative for major differences between these two conditions in the context of active players (**Figure 5D**). For this reason, we focused in revealing active master transcription factors by using a gene regulatory networks (GRN) reconstruction strategy, followed by a modeling of transcription activation cascade, with the help of our previously described tool TETRAMER^20^. This modeling strategy applied to brain organoids trated with BMS753 allowed to infer up to 83 confident master transcription factors (P-value <0.25) presenting a transcription regulation capacity of ∼40 percent; meaning that each of these TFs are able to control 40% of the total nodes composing the GRN (**Figure 5E**). Similarly, 63 confident master transcription factors were inferred from the analysis of the GRN associated to the BMS961+BMS641 treatment (**Figure 5F**). The comparison of the identified master transcription factors for both treatment conditions revealed a set of 45 common players, displayed as part of a gene co-regulatory network, allowing to reveal the transcription regulation relationship between all these factors. (**Figure 5G**).

These master transcription factors were associated to functions like central nervous system development, ventral spinal cord, neural tube development, brain morphogenesis or forebrain generation (**Figure 5H**). Furthermore, several of these players were also associated to specialized neuronal subtypes, retina-related differentiation tissues, as well as mesodermal differentiation giving rise to Microglia (**Figure 5H**). Among the key transcription factors specifically over-expressed under BMS641+BMS961 conditions we have retrieved the Retinal homeobox factor (RAX); known to be expressed in radial glial cells and described as being essential for eye and forebrain development ^21^. Similarly, the expression of the Sine Oculis Homeobox Homolog 2 (SIX2) has been detected; a transcription factor that has been seen to be expressed in the adult differentianting cells of the mouse retina, and from a broader view, in the lens formation in vertebrates ^22^. Furthermore, the transcription factor MAFG has been associated with cataract or lens defects ^23^; the proinflammatory factor BATF2 has been previously shown to be expressed in retinal microglia ^24^; and FOXD1 has been shown to be required for the specification fo the temporal retina ^25^.

Overall the analysis of gexels presenting RAR receptors expression unveiled spatially distinct expression signatures for each of the RAR isotypes, with a rather low fraction of regions presenting their redundant expression. Furthermore, it confirmed the presence of gene expression signatures associated to well-differentiated nervous tissue generation. Notably the use of either the RARα agonist BMS753 or the synergistic activation of the RARβ and RARγ receptors (BMS641+BMS961) gave rise to the activation of a common pool of transcription factors and a subset of others specific to the treatment, and several of the transcription factors specific to the RARβ and RARγ activation appeared to be associated to eye development.

## Discussion

From a historical point of view, developmental processes were studied from a tissue level (morphogenesis), a cellular level (cell specialization) but also from the characterization of molecular components retrieved at precise concentrations in a spatio-temporal fashion (morphogen gradients). Among all these levels of granularity, morphogens were early on shown to be able to control both cell specialization and tissue organization, hence providing a mechanistic view of the events controlling morphogenesis.

Retinoic acid signaling, via the RXR/RAR nuclear receptors activation, is a key morphogen during nervous system development ^1^, but also for brain homeostasis. Indeed, RAR/RXR receptors were shown to be expressed in multiple brain areas in the adult (RARα: hippocampus, cerebellum, cortex; RARβ: striatum, spinal cord; RARγ: hippocampus; RXRα: hippocampus, RXRβ: cerebellum; RXRγ: Striatum, limbic cortex, spinal cord ^26^), and their deletion or a deficiency in retinoic acid supply in mouse models leads to developmental malformations, but also deficits in synaptic plasticity, memory, or accumulation of amyloid-beta in the adult mice ^27–30^.

Despite these findings, brain morphogenesis remains still poorly understood notably from the angle of the gene regulatory programs implicated in its landscaping. Previous studies, including ours, focused in driving neuronal differentiation *in vitro* by the use of the pan-RAR agonist ATRA, but also in decorticating the role of each RAR receptors thanks to the use of specific agonists. Previously we have shown that, in monolayer differentiation assays, mouse stem cells could give rise to neuronal precursors when treated with the RARα specific agonist BMS753, but not in presence of the RARβ agonist BMS641 or the RARγ agonist BMS961 ^6^. Furthermore, we have discovered that the combination of the BMS641&BMS961 agonists can rescue the neuronal precursors differentiation by hijacking the RARα-driven gene programs^7^.

Herein we aimed at addressing first, the influence of these synthetic RAR agonists in long-term monolayer cultures, as a way to interrogate cell heterogeneity during cell specialization; and second, explore their influence in nervous tissue formation.

Cell heterogeneity has been addressed with the help of single-cell transcriptomics assays performed in 16 days monolayer in vitro cultures in presence of either the synergistic action of the BMS641+BMS961 ligands or the RARα specific agonist BMS753. This comparative study revealed a variety of neuronal subtypes retrieved in both treatment conditions, as well as a delayed progression in maturation of cells treated by the combination of BMS641+BMS961. Furthermore, by decoding the chromatin epigenetic status after 16 days of differentiation revealed some differences in promoter activity between both treatment conditions.

Three-dimensional brain organoid cultures during 90 days where the use of Vitamin A has been replaced by either the pan-RAR agonist ATRA, or the use of the combination of BMS641+BMS961 ligands or the RARα specific agonist BMS753, revealed spatially distinct RAR isotype expression, in agreement with the observations in the mouse brain ^26^. Despite a delayed differentiation behavior in presence of the BMS641+BMS961 ligands, the obtained organoids presented gene expression signatures associated to well matured tissues, including a variety of neuronal subtypes but also tissue structure signatures like retina and even the presence of microglia cells.

The presented results are in the front-end of research aiming to generate complex tissues derived *in vitro.* Indeed, while all brain organoid assays rely systematically in the use of Vitamin A as a key component within the maturation medium, the use of RAR-specific ligands remain unexplored. Our results clearly suggest that the use of synthetic agonists can mimic the performance of the classical Vitamin A compound, but it also paves the way for future studies in which further other RAR/RXR-specific ligands can be used for driving tissue specialization.

## Materials and Methods

### Cell Culture

Embryonic stem cells (ESCs) were grown in DMEM supplemented with 4.5 g/l glucose and GlutaMAX-I (11594446; Thermo Fisher Scientific), 15% FBS-ES, 5 ng/ml LIF recombinant mouse protein (15870082; Thermo Fisher Scientific), 1% penicillin–streptomycin, 1% MEM-NEAA, and 0.02% β-mercaptoethanol. For cell differentiation assays, ESCs were seeded onto poly-D-lysine–coated culture plates (0.1%) at a concentration of 100,000 cells per well (for cells collected at 4 days of differentiation), 40,000 cells per well (for cells collected at 8 days treated with ATRA), 20,000 cells per well (for cells collected at 8 days treated with ethanol, BMS753, or BMS961+641; for cells collected at 16 days treated with ATRA), 10,000 cells per well (for cells collected at 16 days treated with ethanol, BMS753, or BMS961+641). The cells are seeded in DMEM (4,5 g/l glucose) w/GLUTAMAX-I, 10% FBS-ES, 1% penicillin–streptomycin, 1% MEM-NEAA, and 0.02% β-mercaptoethanol.

Next day, same medium is refreshed and retinoid ligands are added. ATRA was added for a final concentration of 1 μM. For treatment with RAR subtype–specific agonists, cells were incubated with BMS753 (RARα-specific; 1 μM) or BMS641 + BMS961 (RARβ + RARγ-specific; 0.1 μM each). After 8 days of treatment with either of the aforementioned retinoids, the medium was replaced by Neurobasal medium (11570556; Thermo Fisher Scientific) and B27 without vitamin A (11500446; Thermo Fisher Scientific) and cultured for eight more days. Half medium was refreshed every other day.

### Immunocytochemistry

At the end of the cell differentiation assay, medium is aspirated completely and cells are fixed in 4% paraformaldehyde followed by 3 × 5 min washes in PBS. Cells were permeabilized (Triton 0.1% in PBS; 15 min at room temperature), washed 3 × 5 min in PBS and blocked (0.1% Triton and 1% BSA in PBS for 1 h at room temperature). Cells were incubated with the primary antibodies overnight at +4°C [anti-β III tubulin/ anti-TUBB3 (ab14545; Abcam), anti-MAP2 (ab32454)]. Cells are washed 3 × 5 min in PBS followed by incubation with a secondary antibody for 1 h at room temperature [(donkey anti-mouse IgG [H + L] antibody Alexa 555 (Invitrogen A-31570), donkey anti-rabbit IgG [H + L] antibody Alexa 488 (Invitrogen A-21206)] respectively. Cells are washed 3 × 5 min in PBS and finally mounted on microscope slides with mounting medium containing DAPI (P36962; Thermo Fisher Scientific).

### RNA extraction and cDNA generation

Total RNA was extracted using RNeasy Mini Kit (74104; Qiagen). 0.5 – 1 µg of RNA was used for reverse transcription (4368814; Applied Biosystems). Transcribed cDNA was diluted fivefold and used for real-time quantitative PCR with QuantiTect SYBR Green PCR Kit (204145; Qiagen). Comparative Ct (ΔΔCt) method was used for performing relative quantitation of gene expression.

### Single cell RNA sequencing and processing

The Chromium instrument and Single cell 3’ Reagent Kit v3 (PN-1000128; 10X Genomics) was used to prepare individually barcoded single-cell RNA-Seq whole transcriptome libraries following the manufacturer’s protocol. The target number of cells was 8000 per condition. Libraries were sequenced on Illumina Novaseq 6000 using paired-end 150 cycles v3 kits (Read 1: 28 cycles; Index Read 1 (i7): 10 cycles; Read 2: 90 cycles).

### Single cell transcriptome

Data from 9 conditions has been processed using Cell Ranger 7.0.1 for primary analyses, aligned to the mouse reference genome mm10 2020A. Cells were filtered out on the grounds of doublets (DoubletFinder V.2)(McGinnis et al., 2019), more than 5% of mitochondrial counts content, novelty score higher than 0.8, and cells with less than 200 genes. These cells from different conditions were merged and integrated by a selected group of genes (200 features). Read counts were normalized by SCTransform method and processed downstream for Uniform Manifold Approximation and Projection (UMAP). Clusters were identified (KNN method) based on gene expression similarities. Differential expression analysis was performed and filtered out genes with a log2 fold change < 0.25 and a p value > 0.05 using the Wilcoxon test. All these steps were performed in R (R core team 2024) using the package “Seurat” (Version 5) (Hao et al., 2024) and customized scripts.

For Figure 1E, the fraction of cells per sample assigned to each cluster was calculated and further normalized to the total levels per time-point. These frequencies were clustered by Euclidian distance and displayed in a heatmap (MeV: Multiple Experiment Viewer). In the Figure 1F a P-value in a negative logarithmic scale in base 10, scaled 10x was calculated by a hypergeometric test from the differential expression analysis and the target gene sets, using a customized “R” script. These “gene set” collections were manually curated from different publications or databases (Supplementary table S2).

Pseudo time analysis was performed using the python package “STREAM” (Chen et al., 2019) in all the 9 samples together (3 treatments, 3 time points). To perform a dimension reduction, the next parameters were set up: 2,000 variable genes were selected, lineal method “mlle”, 10 principal components and 50 neighbors. The trajectory was inferred by the dimension manifold projection (UMAP). The next parameters were used to tunning the trajectory: epg_alpha=0.02, epg_mu=0.1, epg_lambda=0.02, and epg_ext_par=0.8 for the branch extension. A flat tree was obtained with a dist_scale=0.5. Ribbon rooted trees were graphed with the parameter distance scale=2 showing the 3 treatments together (Fig. 2A), or the different timepoints (Fig. 2B). Percentage of cells contained in each branch by different treatment conditions, or compared to by different clusters from in the UMAP projection in Fig. 1 C were calculated, depicted and clustered (method “complete”) in a heatmap (Fig. 2 C and D).

A hypergeometric test to obtain the probability of finding the highly expressed genes from each branch (showed in the figure 2A) in different gene sets was performed. The fold change was calculated dividing the mean value for determinate gene and branch, over the total mean counts for the determinate gene. Genes with fold change > 0.5 in logarithmic values on base 2 were selected. The P-values obtained were transformed to a negative logarithmic scale on base 10, scaled 10x, clustered and depicted in figure 2E using a customized R script. Details from these gene sets can be found in Supplementary table S2.

### Cut&tag epigenomics assays

Cut&Tag experiments were carried out as previously described (Kaya-Okur et al^11^; Li et al^31^), with some modifications. Briefly, 250.000 cells were captured with BioMagPlus Concanavalin A (Polysciences, 86057-3) and incubated with the appropriate primary antibody overnight at 4°C (anti-Histone H3 (tri methyl K4) antibody (Abcam, ab8580; 1:50); anti-Histone H3(acetyl K27) antibody (Abcam, ab4729; 1:50); anti-Histone H3(tri methyl K27) antibody (Abcam, ab192985; 1:50)); Anti- RNA Pol II monoclonal antibody: diagenode (C15100055) (1:20). For the experiment in which the antibody RNA Pol II was used, the mixture cells-beads were fixed with 1% PFA in 1xPBS for 15 minutes before the addition of the primary antibody. A secondary antibody was incubated at room temperature for 1 hour (Donkey anti-rabbit IgG antibody: Sigma-Aldrich, SAB3700932; 1:100). Fusion protein ProteinA-Tn5 was prepared in-house (plasmid Addgene #124601) by following a previously described protocol (Li et al. ^31^). Purified PA-Tn5 was loaded with the following adapters compatible with Illumina sequencing (hybridized with the complementary Mosaic sequence [PHO]CTGTCTCTTATACACATCT): MOS-universal: TACACTCTTTCCCTACACGACGCTCTTCCGATCTAGATGTGTATAAGAGACAG MOS-index: GTTCAGACGTGT.GCTCTTCCGATCTAGATGTGTATAAGAGACAG Loaded PA-Tn5 complex (0.16 μM) was incubated for 1 hour at RT, followed by tagmentation step for 2 hours at 37°C. DNA was extracted by phenol/chloroform (Sigma-Aldrich 77617) followed by ethanol precipitation. Finally, the tagmented DNA was amplified using Illumina primers (NEBNext® Multiplex Oligos for Illumina, NEB #7335S) with the following cycling conditions: 98°C for 30s, 20 cycles of 98°C for 10s and 65°C for 75 s; then final extension at 65°C for 5 min and hold at 12°C. DNA libraries were cleaned using 1x volume of SPRI-select beads (Beckman Coulter, B23318) and Illumina sequenced (150-nt pair-end sequencing; Illumina NovaSeq 6000).

Cut&tag histone modification sequencing data were aligned to the mouse genome (mm9) using Bowtie2 (v2.4.1) under the default parameters. Chromatin state analysis has been performed with ChromHMM (v1.14). Bedgraph files and peak annotations were obtained with MACS2 and visualized within the IGV genome browser (v.2.4.15).

### Mouse brain organoid

Mouse brain organoid protocol has been derived from Lancaster et al, 2013. ESCs should be present in culture as EBs (embryoid bodies). On Day 0, the EBs are dissociated with accutase (A6964-100ML; Sigma) for 10 min. Cells are seeded in P96 (ultra low attachment) plate at 5000 cells/ well in 150µl medium/ well [DMEM (4,5 g/l glucose) w/GLUTAMAX-I, 10% FBS-ES, 1% penicillin–streptomycin, 1% MEM-NEAA, and 0.02% β-mercaptoethanol, SB 431542 (1/1000)] and incubated for 2 days at 37°C. On Day 2, same medium is refreshed by taking out 75µl and adding 150µl fresh medium per well. On Day 4, medium is refreshed by taking out 150µl medium and adding 170µl Neural induction medium/ well (DMEM/F12, 1% N2 supplement, 1% Glutamax supplement, 1% MEM-NEAA, 1% Penicilline –Streptomycine, 0.1% Heparin solution). On day 6, the EBs are embedded in Matrigel GF reduced (11523550; Fisher scientific) for 45 min before transferring to P6 (ultra low attachment) plates in maturation medium (DMEM/F12 medium, Neurobasal medium, 2% B27 without vitamin A supplement, 1% Glutamax supplement, 1% Penicilline –Streptomycine, 0.5% N2 supplement, 0.5% MEM-NEAA, 0.025% insulin (I9278-5ML; Sigma). On day 9, medium is changed completely to maturation medium and adding 2% B27 with vitamin A supplement (11530536; Thermo Fisher Scientific) or retinoid ligands for a final concentration of [(ATRA (1µM), BMS753 (1µM); BMS641, BMS961, BMS641+961 (0.1µM)] and placed on an orbital shaker at 75-80 rpm. Two-thirds of medium is changed twice a week.

### Brain organoids cryo-sectioning and spatial transcriptomics library preparation

Organoids are fixed with 4% paraformaldehyde, 6% sucrose for 20 min, washed 3 x 5 min with PBS and suspended in 30% sucrose at 4°C overnight. Several organoids are placed in plastic molds and submerged in OCT compound and frozen before being cryo-sectioned using a cryo-star. Frozen brain organoid molds are cryosectioned and deposited on top of in-house manufactured DNA arrays as described previously (Lozachmeur et al. ^18^). Briefly, in-house manufactured DNA arrays are composed by a double-barcoded part encoding their unique position within the array, and a poly-T extremity allowing to capture messanger RNA.

During organoids cryosectioning, slides were kept inside the cryostat chamber in order to conserve the integrity of the RNA and at −80°C for long storage. Deposited tissue sections were processed for library preparation as described in our in-house protocol (Lozachmeur et al.^18^). At the end of the process, libraries were used for Illumina sequencing (150 nts paired-ends sequencing; NovaSeq; >200 million reads per library).

### Spatial transcriptomics processing

Primary analysis has been performed with our in-house developped tool SysISTD (SysFate Illumina Spatial transcriptomics Demultiplexer: https://github.com/SysFate/SysISTD). SysISTD generates as outcome a matrix presenting read counts per gene IDs in rows and spatial coordinates in columns. To focus the downstream analysis to the physical positions corresponding to the analyzed tissue, we used an in-house R script taking as entry an image of the DNA array scanned with the TRICT filter, releaving the presence of the fiducial borders introduced during the manufacturing of the DNA arrays.

The focused matrix was quantile normalized with our previously described tool MULTILAYER^32,33^. Then, the normalized matrix was further processed for revealing cell/tissue density contrast information by integrating normalized counts with Gexel (gene expression element, in analogy to pixel) value levels retrieved in the TRITC tissue image. The obtained normalized and pixels intensity adjusted matrix was used for all downstream analysis.

Gexels classification were performed by processing the normalized read count matrices to obtain the pseudo-time information using the python package “STREAM” (Chen et al., 2019 ^10^), in the same way as the single cell data (described above) except for the next parameters: For dimension reduction a loess_frac=0.1, with a previous normalization by ‘lib_size’, two principal components and 15 neighbors, to tunning the trajectory: epg_alpha=0.01, epg_lambda=0.08, epg_trimmingradius=0.1, and the branching was optimized with incr_n_nodes=30. Before this step, gexels with total number of counts lower than the first quartile (0.25) of the total read counts in the matrix were filtered out.

Gexels presenting upregulated levels for each retinoid receptor (Rara, Rarb, or Rarg) across treatment samples were selected using the second quartile of the normalized counts from all pixels for each respective gene. The pixels with lower number of counts were filtered out like indicated before.

### Gene regulatory network reconstruction

Highly expressed genes for each condition (except for the intersection of downregulated) were analyzed with our previous described tool TETRAMER (Cholley et al. ^20^) within the “Cytoscape” platform. Briefly, a gene regulatory network (GRN) was generated by using a collection of Transcription Factor-Target gene relationships (CellNet database; available within TETRAMER), which has been integrated with the differentially expressed genes retrieved within the Gexels presenting high levels of Rar receptors.

TETRAMER inferred a master regulator index for each TF retrieved in the network, which were used for reconstructing a gene co-regulatory view among the top ranked factors. The betweenness centrality was calculated for this gene co-regulatory network considering the direction of the edges, this topological parameter provides the centralized structure to the network. Cell/tissue annotations for genes composing this gene co-regulatory network has been performed with GeneMania^34^. Obtained gene associations to the cell/tissue annotations were merged with cell type annotations obtained from Supplementary Table S2, and displayed in a binary heatmap format with MEV.

## Data access

All datasets generated on this study have been submitted to the NCBI Gene Expression Omnibus (GEO; http://www.ncbi.nlm.nih.gov/geo/) under accession number GSE XXX.

## Acknowledgements

We thank all members of the <u>team SysFate</u> for contributing to the discussion of this project and the Genoscope sequencing platform for their technical support. Further acknowledgments to Celine Derbois from the platform of the CNRGH, for the help in the preparation of scRNA-seq libraries, and to Gwendoline Lozachmeur from SysFate for the help in the preparation of spatial transcriptomics libraries. This work was supported by the institutional bodies CEA, CNRS, Université d’Evry-Val d’Essonne. A. K has been supported by the “Fondation pour la recherche medicale” (FRM; funding ALZ°201912009904); A. G is supported by EU grant HORIZON N° 101070740; M.G.M.F has been funded by the Institut National du Cancer (INCa: Funding N° 2020-124; N° 2022-078).

## Author contributions

A Galindo-Albarra: investigation, data curation, software, and formal analysis. A. Koshy: formal analysis, investigation, and methodology. MG Mendoza-Ferri: resources and methodology. MA Mendoza-Parra: conceptualization, formal analysis, supervision, funding acquisition, and writing—original draft.

## Conflict of interest

The authors declare that they have no conflict of interest.

